# Calculating the Human Mutation Rate by Using a NUMT from the Early Oligocene

**DOI:** 10.1101/016428

**Authors:** Ian Logan

## Abstract

As the number of whole genomes available for study increases, so also does the opportunity to find unsuspected features hidden within our genetic code. One such feature allows for an estimate of the Human Mutation Rate in human chromosomes to be made. A NUMT is a small fragment of the mitochondrial DNA that enters the nucleus of a cell, gets captured by a chromosome and thereafter passed on from generation to generation. Over the millions of years of evolution, this unexpected phenomenon has happened many times. But it is usually very difficult to be able to say just when a NUMT might have been created. However, this paper presents evidence to show that for one particular NUMT the date of formation was around 29 million ago, which places the event in the Early Oligocene; when our ancestors were small monkey-like creatures. So now all of us carry this NUMT in each of our cells as do Old World Monkeys, the Great Apes and our nearest relations, the Chimpanzees. The estimate of the Human Mutation obtained by the method outlined here gives a value which is higher than has been generally found; but this new value perhaps only applies to non-coding regions of the Human genome where there is little, if any, selection pressure against new mutations.

## INTRODUCTION

In the last few years an increasing number of primate species have had their genomes sequenced and the results placed in the Public Domain. The first genome was produced by the Human Genome Project, and now there are genomes for 15 species of Primate available for study at http://www.ncbi.nlm.nih.gov/mapview/

The activity of examining a genome for a particular purpose has been described as data mining (Yao 2009); and there is no doubt that many interesting features of our DNA are yet to be found. The human genome contains the *coding* for about 20,000 active genes and this accounts for under a half of the DNA in the chromosomes, with the remainder being *non-coding*; and it is the data mining of the *non-coding* regions that is likely to yield the greatest surprises. In this paper a particular *non-coding* region is examined with the purpose of calculating the Human Mutation Rate.

Lipson et al. (2015) in their recent paper reviewed the importance of calculating the Human Mutation Rate and suggested 4 methods that can be used in its determination. The methods were – described in simple terms: [1] comparing corresponding DNA sequences DNA from different modern day species and counting the mutations that have occurred over an estimated period of time; [2] comparing genomes from related modern day people and counting the observed differences; [3] comparing a modern day genome to an archaic genome, for example from a Neandertal or a Denisovan; and [4] a mathematical method termed *Ancestral recombination Density*. These 4 methods give a Human Mutation rate (*µ*) in the region of: *µ = 1-2.5 x 10*^*−8*^ *per base per generation* (using 29 years as a generation)

But what does this really mean ? And, a simpler concept might be to consider the percentage change in a DNA sequence, making *µ* as described above equivalent to; On average, *1-2.5% of the bases mutate in a period of 29 million years*

This Human Mutation Rate is assumed to apply to the total DNA of the chromosomes, however *coding* DNA can be assumed to mutate at a much slower rate than *non-coding* DNA because of *selection* - with harmful mutations being *selected* against. Whereas, the mutation rate in a *non-coding* sequence has no such restriction and the mutation rate will be much higher.

In this paper a *non-coding* sequence, a mitochondrial NUMT, is described and used to calculate the Human Mutation rate as it applies to Chromosome 8; with the result being that a value for *µ* is found that is higher than previously described.

## Mitochondrial DNA

In every cell of the body as it is formed there is a nucleus, containing 23 pairs of chromosomes, and in the cytoplasm are the mitochondria. These are small structures associated with the production and transfer of energy within the cell and they contain short DNA molecules– the mitochondrial DNA (mtDNA). These molecules contain about 16,569 bases and in a person most of mtDNA molecules will be identical. The bases of the mtDNA can be considered as being numbered and their state (A, C, G or T) is usually described compared to the Cambridge Reference Sequence (CRS). (Anderson 1981, Andrews 1999)

The bases in the mtDNA do have a high mutation rate, much greater than that of the chromosomal DNA, with roughly 50 mutations having occurred in the last 200,000 years. Mutations can occur in all parts of the mtDNA, but there is *selection* against overtly harmful mutations being propagated, with the result that about 5 times as many mutations are observed in *non-coding* areas as opposed to *coding* areas. (Logan, 2007, Behar 2012)

The structure of the mtDNA has been extensively studied and in each molecule there is *coding* for 13 genes, 2 RNA molecules, and 22 tRNAs and sequences associated with the production of further mitochondria – such as the ‘OriL’ sequence concerned with the replication of the *light* strand. Also, and of importance in respect to this paper, are a number of *non-coding* sequences interspersed between the functional *coding* sequences.

In this paper the area of the mtDNA from base 5097 to base 7195 is of special interest and corresponds to: 2 incomplete genes (MT-ND2 & MT-CO1), 5 tRNAs (for the amino acids Tryptophan, Alanine, Asparagine, Cysteine & Tyrosine), the OriL region, and 3 *non-coding* regions. Table 1 shows this area of the mtDNA in some detail.

**Table 1:**
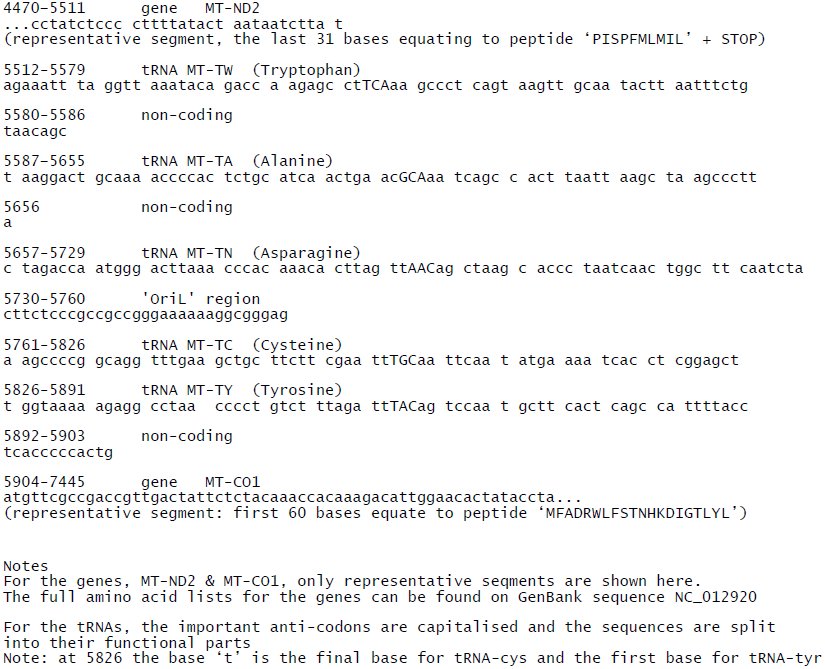
The parts of the mtDNA (CRS) for the region of interest (bases 5097-7195)

## NUMT (NUclear MiTochondrial DNA)

A NUMT (pronounced as ‘new-might’) is a fragment of the mitochondrial DNA (mtDNA) that has become incorporated into a chromosome. Initially, it was thought this type of *capture* was an uncommon occurrence, but now it is suspected that the formation of NUMTs has occurred in all species and, although infrequent, is an ongoing process. (Selected references: Herrnstadt 2009, Bensasson 2003, Hazkani-Covo 2008, Logan 2009, Simone 2011, Calabrese 2012)

Recently, a paper detailed a number of NUMTs that are to be found in the people from different populations around the world, showing that the formation of these NUMTs date to less than 200 thousand years. (Dayama 2014) By the very nature of the method of its formation, a NUMT shows several random features:

Firstly, the fragment separated from the mtDNA can be small or large. However, in most instances it is not possible to data mine for small NUMTs, perhaps of less than 50 bases in length, but with increasing size the searching becomes easier. Now, over a thousand NUMTs have been identified in the Human genome.

Secondly, the actual chromosome *capturing* a NUMT appears to another random event. However, the number of NUMTs in a given chromosome is closely related to the overall size of the chromosome, with the largest, chromosome 1, having a considerable number of NUMTs.

Thirdly, the position of a NUMT within a chromosome again appears to be essentially random. However, there are reports describing NUMTs that have interfered with gene *coding* regions and caused disease. (Reviewed by Hazkani-Covo 2010)

Fourthly, the bases in a NUMT can themselves mutate over the generations, and as with any other mutation in a *non-coding* region there is unlikely to be *selection* against a mutation in a NUMT.

Fifthly, there is the question of *age determination*. A NUMT that has a sequence that closely matches a modern mtDNA sequence can be considered to be of recent origin. So if a NUMT shows almost total concordance it is likely to have been formed within, say, the last 5 million years. But if the match is as poor as 75% concordance, such a NUMT might have been formed, say, 30 million years ago. Unfortunately, it is not possible to ascertain the exact age of a given NUMT and there will always be a degree of uncertainty over any age that is suggested.

And, finally, there is the feature of inter-species *conservation*. A Human NUMT with an age of formation of more than, say, 6 million years is likely to be found in the genome of our closest relative, the Chimpanzee. And, with a Human NUMT formed, say, 30 million years ago, it might expected to be found in all the genomes of the Old World Monkeys, the Great Apes, and, of course, Homo sapiens.

## The Primate Evolutionary Tree (See Figure 1)

The exact dates involved in the Primate Evolutionary tree are uncertain, but the details in Figure 1 are probably reasonable. (Finstermeier 2013, Pozzi 2014)

**Figure1:**
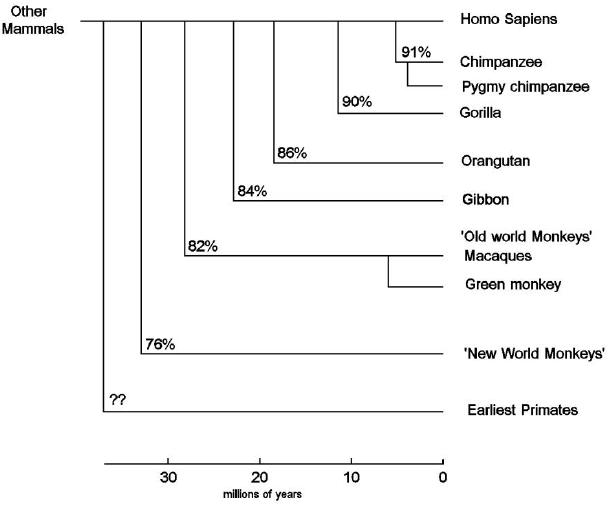
The Primate Evolutionary Tree The horizontal scale is measured in millions of years. The percentages show the mtDNA relatedness.

To summarise, the figure shows modern day man as having separated from:

- the Chimpanzee and Bonobo around 6 million years ago
- the Great Apes (i.e. the Gorilla, Orangutan, and Gibbon) around 20 million years ago
- the Old World Monkeys (i.e. the Macaques and the Green monkey) around 25-30 million years ago.
- and the New World Monkeys and the earliest Primates (i.e. the Golden snub-nosed monkey, the Bolivian Squirrel Monkey, the White-tufted-ear marmoset, the Philippine tarsier and the Small-eared galago) more than 30 million years ago.

In this paper the period of 29 million years will be used to describe the separation between modern day man and the Green monkey solely because this figure allows the separation to be considered as having occurred 1 million generations ago.

## Materials and methods

This paper identifies a selected region of Chromosome 8 as being a NUMT; and uses 4 DNA sequences as evidence to support this hypothesis. The sequences are all obtainable from the NCBI’s GenBank database.

The first sequence shows the region of the Human mtDNA sequence, the CRS, from bases 5097-7195. Details of how to obtain this sequence, and the 3 other sequences, is given in a step-by-step guide as part of the supplementary material.

The parts of this region of the first sequence are listed in table 1.

The second sequence is the presumed NUMT, from chromosome 8, coordinates 110933244-110935358 This NUMT has been also classified as HSA_NumtS_321 by Calabrese (2012). It is to be noted that there are several NUMTs in the Human genome that match the CRS region 5097-7195. But there does appear to be only a single NUMT from chromosome 8.

The third and fourth sequences are the corresponding mtDNA and NUMT sequences from the genome of the Green monkey (Chlorocebus sabaeus).

For the purposes of this paper the genome of the Green monkey has been chosen, and the appropriate details provided, because it is the genome of a species which is the most distant in evolutionary terms from Homo sapiens and yet still shows the same NUMT. The genomes of the Rhesus macaque or the Crab-eating macaque would serve just as well.

Also, for clarity, a few bases at ends of the NUMT have not been selected as the bases do appear to vary between one genome and another.

## Results

The hypothesis suggested in this paper is firstly, that a NUMT corresponding to the mtDNA region 5097-7195 exists and can be found on Chromosome 8, and secondly it is suggested that this NUMT was formed about 29 million years ago. And, In order to support the two parts of the hypothesis 4 DNA sequences have been obtained from the GenBank database.

The first sequence comes from the mitochondrial genome of the Human, as represented by the CRS. The sequence contains the *coding* for parts of 2 genes, 5 tRNAs, the ‘Oril’ region, together with 3 *non-coding* regions.

The second sequence is from a region of Chromosome 8 and this is shown to be a NUMT as the sequence can be mapped against the first sequence without difficulty.

Table 2 shows the mapping of the CRS sequence against the NUMT and shows that all the different functional parts present in the CRS sequence can be demonstrated in the NUMT.

**Table 2:**
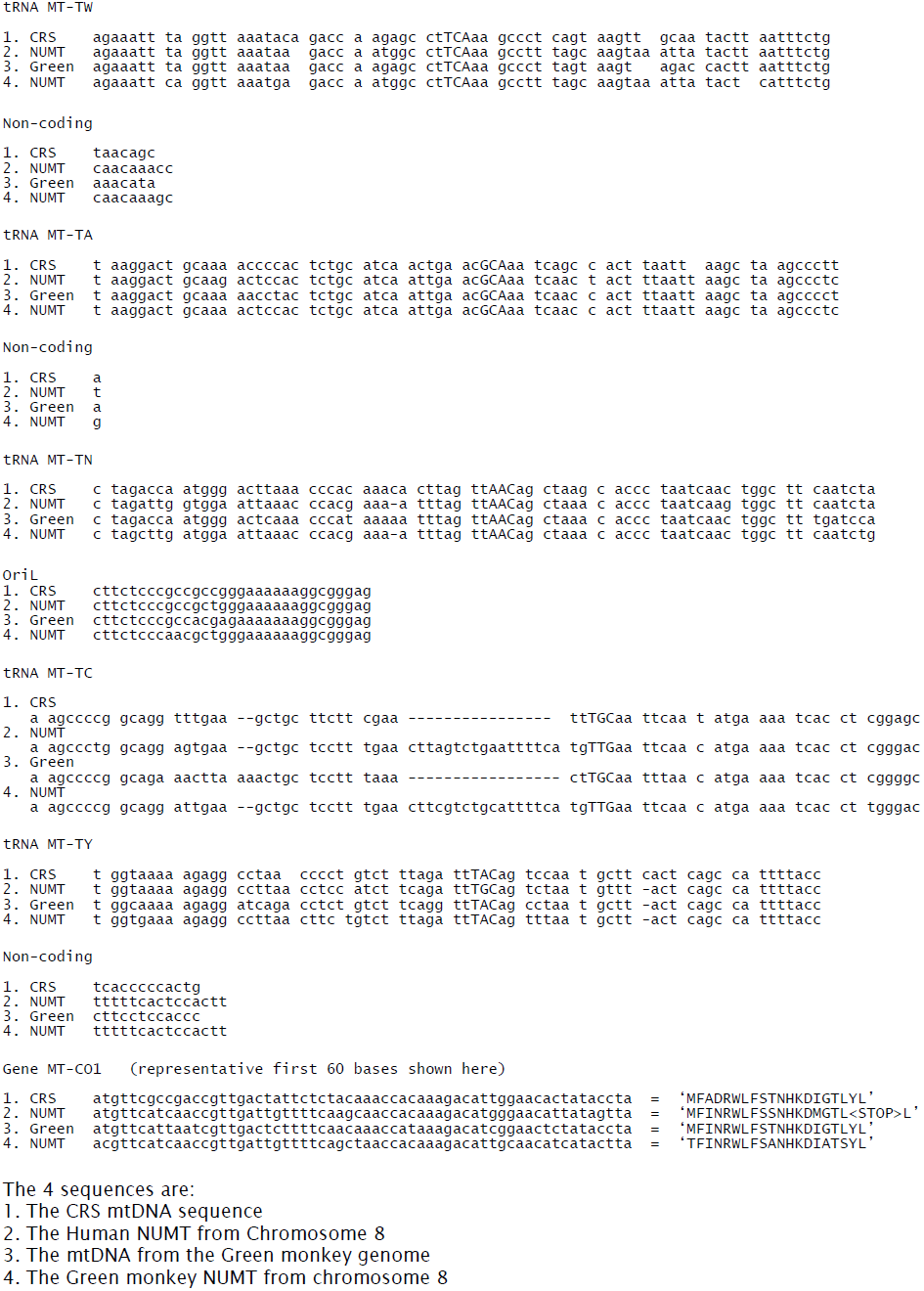
The 4 sequences compared

The third and fourth sequences repeat the above process, but this time using sequences from the genome of the Green Monkey.

Once again Table 2 shows how the 4 sequences map against each other in respect of all the different functional parts. It is interesting to note that both the NUMT from the Human genome and that from the genome of the Green monkey have the same *insertion* of 16 bases in the tRNA sequence for the amino acid, Cysteine.

The presence of NUMTs that match each other and are found in the genomes of 2 species of primate that are considered to have separated from each other around 29 million years ago, does suggest that this NUMT was indeed formed at the time of separation, and in the Early Oligocene.

Whilst all the sequences map against each other, there are mutational differences between the sequences and the number of observable mutations is crucial to the estimation of the Human Mutation Rate.

And, http://www.ncbi.nlm.nih.gov/mapview/ and BLAST (Altschul 1990) can be used to show that the correspondence between the Human NUMT sequence and that of the modern Human mtDNA sequence is about 76%, or putting it another way, about 24% of the bases of the NUMT show mutations. However, this method of comparing a NUMT sequence from a chromosome with a modern mtDNA sequence gives a very high figure as the mutation rate in mitochondrial DNA is very much higher than in chromosomal DNA.

But it is possible to calculate the Human Mutational Rate by considering the mutational changes that show as differences in the DNA sequences of the two NUMTs from the Human and the Green monkey. This method looks for the mutations that have occurred over a period of 29 million years in both species and therefore half the number of differences can be assumed to have occurred in just one NUMT.

The number of differences between the 2 NUMTs is approximately 7.6 % (150 base mutations and 14 small insertions/deletions in the 2,115 bases) which suggests a Mutation Rate value of *µ = 3.8 x 10*^*−8*^ per base per generation.

## Discussion

The claim to being able to calculate the Human Mutation Rate by using a new method has to be justified carefully.

In this study the existence of a NUMT corresponding to a substantial part of the mtDNA molecule has been shown. The NUMT found on Chromosome 8 is over 2,100 bases in length and maps against the mtDNA sequence of modern day man, using the CRS as the example, in respect of gene *coding*, multiple tRNA *coding,* the special OriL region, as well as several distinctively sized *non-coding* regions. Repeating the process with the genome of our distant relative the Green monkey also shows that the same NUMT is to be found on its chromosome 8. This NUMT from a distantly related species also shows the same distinctive features, which strongly suggest the common origin of both NUMTs in a single event; and the subsequent conservation in the two species.

The method therefore can be considered as showing that the original NUMT was formed around 29 million years ago. (This figure used as it represents 1 million generations.) But there has to be considerable uncertainty over this figure and even to use the wide range of +/- 5 million years might not be unreasonable.

Also, there is the question of whether it is reasonable to compare the NUMT from the modern Human mtDNA sequence against the NUMT from the Green monkey to obtain a measure of the Human Mutation Rate. Clearly, bases within the NUMT will have mutated since its formation, and it appears that perhaps 7.6% of them have perhaps done so. This figure shows the mutations that have occurred in both NUMTs and therefore the figure has to be halved to obtain the Mutation Rate as it applies just to the Human.

For the above reasons, a suggested value of *µ = 3.8 x 10*^*-8*^ per base per generation must be considered as having a wide margin of uncertainty, although it is hard to believe the rate would be higher. Therefore, it is perhaps best to consider that this new method of calculating the Human Mutation Rate gives a value of *µ = 3 - 4 x 10*^*−8*^ per base per generation. This value is higher than previous estimates, and probably reflects that in purely *non-coding* DNA the mutation rate is high when there is little, if any, selection against mutations.

The paper tries to highlight the difficulties in obtaining a meaningful figure for the Human Mutation Rate when it is not possible to compare DNA sequences directly. The use of a NUMT, which is considered to a be ‘fossilised’ record of the mtDNA as it was at the time of the formation of the NUMT is an attempt to obtain an archaic sequence which can then be compared to a modern day sequence. This method does perhaps resolve some of the problems used in other methods (as reviewed by Lipson 2015).

Also, there is the question as to whether the term ‘Human Mutation Rate’ can be considered to refer to a single quantity. Or, is there one figure for *coding* regions, another for *non-coding* regions, and a third overall figure. In this paper the suggested Human Mutation Rate applies only to *non-coding* regions.

And, is it reasonable to use a figure of 29 years per generation when considering mutations that have occurred since the Oligocene in the Human ancestral line and that of the Green monkey.

Overall the method does give a useful value for the Human Mutation Rate that can be compared to that obtained by other methods.

## Supplementary material

There are 2 parts to the supplementary material

Supplement: A A step-by-step guide showing how to obtain the 4 sequences discussed in the paper.

Supplement: B The 4 sequences identified in the paper given in FASTA format

### Supplement A: A step-by-step guide showing how to obtain the 4 sequences discussed in the paper

In order to keep the paper as simple as possible, the selection of sequences is restricted to just 4 sequences – all obtainable from the NCBI GenBank database.

The first sequence: Human mtDNA (CRS)

By using: http://www.ncbi.nlm.nih.gov/nuccore/NC_012920 With the *Change region shown* set *from:* 5097 & *to:* 7195 Click on *Update View*

This gives sequence of 2099 bases from the Human mtDNA (CRS).

The second sequence: The Human NUMT

Open http://www.ncbi.nlm.nih.gov/mapview/

Following along the Homo sapiens line click on B (for BLAST) Enter the first sequence into the box

Select *Somewhat similar sequences (blastn)*

Select BLAST

When the hits appear, select the NUMT for Chromosome 8 This shows the NUMT is to be found on sequence NC_000008 coordinates 110933244-110935358

And can be fetched from: http://www.ncbi.nlm.nih.gov/nuccore/NC_000008 Use *Customize view* & *show reverse complement* to get the matching orientation.

This gives the corresponding Human NUMT.

The third sequence: The Green Monkey mtDNA Open http://www.ncbi.nlm.nih.gov/mapview/

Following along the line Chlorocebus sabaeus click on B (for BLAST) And using the first sequence and *blastn*,

Shows the corresponding Green Monkey mtDNA sequence is located at: NC_008066 5092-7190

And this sequence can be downloaded from: http://www.ncbi.nlm.nih.gov/nuccore/NC_008066

The fourth sequence: The Green Monkey NUMT from Chromosome 8

This is obtainable from: http://www.ncbi.nlm.nih.gov/mapview/

Following along the line Chlorocebus sabaeus click on B (for BLAST) and inputting the second sequence and using *blastn.*

The NUMT is found on NC_023649 at 105641836-105643957

And the sequence can be fetched from: http://www.ncbi.nlm.nih.gov/nuccore/NC_023649

using *from:* 105641836 & *to:* 105643957 and *show reverse complement*

### Supplement: b: The 4 sequences identified in the text listed as FASTA format

1. The 2099 base sequence from the Human mtDNA 5097-7195 NC_012920

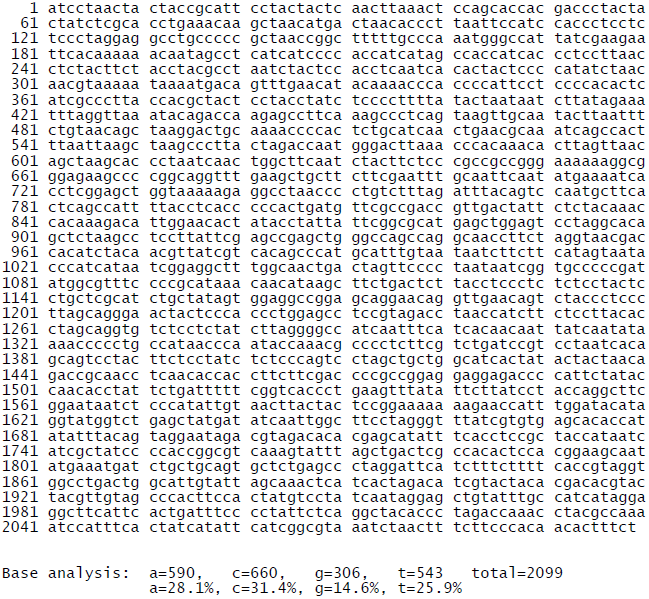

2. The Human Chromosome 8 NUMT

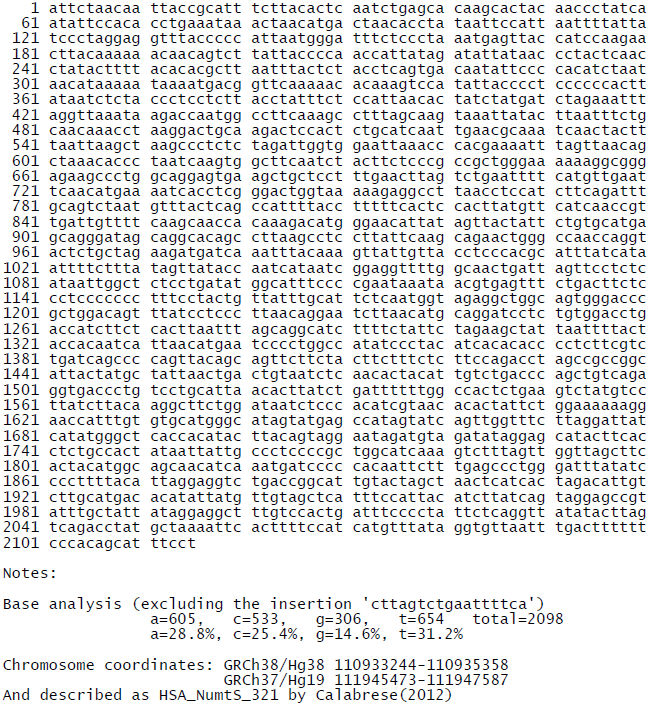

3. Green Monkey mtDNA Bases 5092-7190

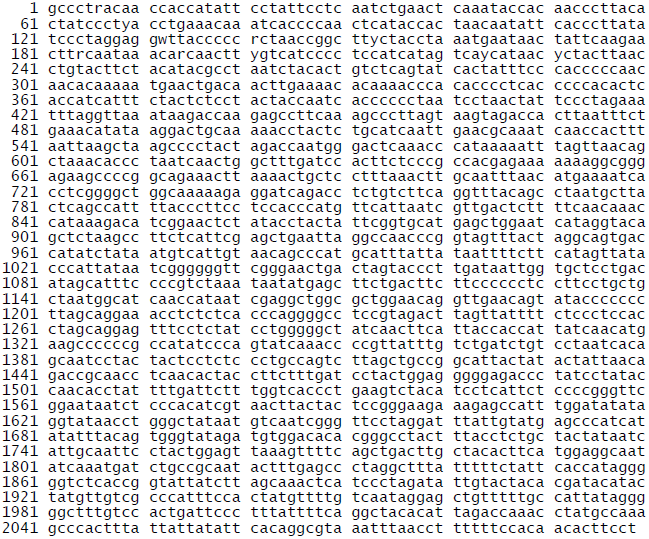

4. Green Monkey NUMT on Chromosome 8

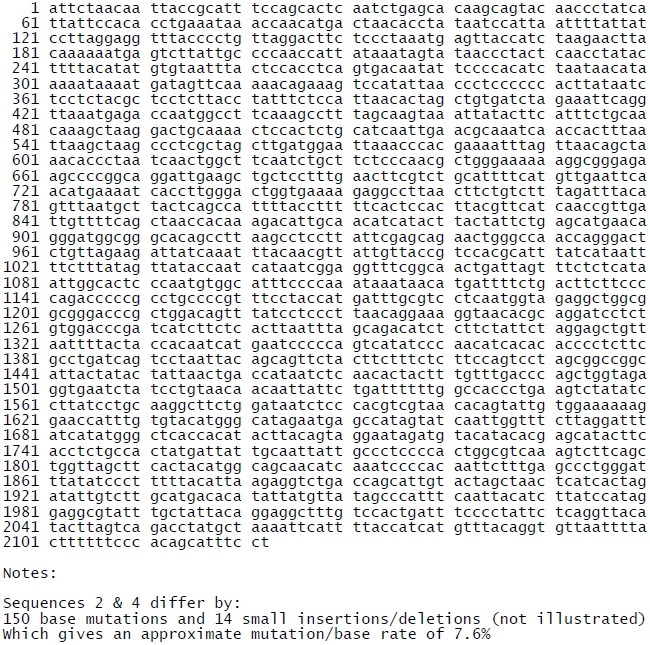

